# TET1 Functions as a Tumor Suppressor in Lung Adenocarcinoma Through Epigenetic Remodeling and Immune Modulation

**DOI:** 10.1101/2024.10.31.621362

**Authors:** Abdur Rahim, Brian Ruis, Andrew T. Rajczewski, Monica Kruk, Natalia Y. Tretyakova

## Abstract

Ten-Eleven Translocation (TET1-3) dioxygenases oxidize 5-methylcytosine (5mC) in DNA to generate 5-hydroxymethylcytosine (5hmC), 5-formylcytosine (5fC), and 5-carboxylcytosine (5caC), initiating DNA demethylation. The three proteins share significant sequence homology and catalyze the same chemical reaction utilizing alpha-ketoglutarate cofactor and non-heme iron to oxidize the methyl group of 5mC. Since their discovery in 2009, there have been contradictory reports regarding the roles of TET proteins in cancer. *TET* genes have been characterized as tumor suppressor genes because their expression levels are reduced in many human cancers including lymphoma, prostate, and pancreas, and *TET2* gene mutations are common in hematological cancers. However, *TET1* was recently reported to be overexpressed in triple negative breast cancer and to act as a protooncogene in lung cancer. In the present study, we employed genetic approaches to directly address the function of TET1 protein in lung adenocarcinoma. We found that overexpression of *TET1* in human lung adenocarcinoma (H441) cells decreased their proliferation and inhibited colony formation, cell migration, and 3D spheroid tumorigenesis. In contrast, *TET1* knockout in lung adenocarcinoma accelerated cell growth and promoted colony formation, cell migration, and 3D spheroid tumorigenesis. Transcriptomics and proteomics analyses revealed that *TET1* overexpression was associated with overexpression of immune markers, primarily via activation of TNF and NF-kB signaling pathways. *TET1* knockout in lung adenocarcinoma cells induces the expression of genes involved in cellular metabolism and cell growth. Our results are consistent with a tumor suppressor role of *TET1* gene in lung adenocarcinoma and reveal its role in activating antitumor immunity.

## Introduction

DNA methylation (5-methylcytosine) is a major mechanism by which eukaryotic cells maintain tissue-specific patterns of gene expression.^[1]^ Cytosine methylation takes place primarily in the context of CpG dinucleotides and is catalyzed by DNA methyltransferases (DNMT).^[2]^ DNA methylation marks are introduced by *de novo* methylases DNMT3a and DNMT3b and are maintained during semiconservative replication by DNMT1.^[3]^ CpG methylation upstream of transcriptional start sites is typically associated with reduced levels of gene expression,^[4]^ while cytosine methylation within gene bodies may have an opposite effect.^[5]^ The presence of the C-5 methyl group on cytosine mediates binding of methyl-CpG binding proteins to promoter sequences, followed by the recruitment of histone-modifying enzymes and the formation of closed chromatin.^[6, 7]^ Ten eleven translocation (TET1-3) dioxygenases utilize non-heme iron and α-ketoglutarate (α KG) to catalyze the oxidation of (5mC) in DNA to 5-hydroxymethyl-cytosine (5hmC), 5-formylcytosine (5fC), and 5-carboxylcytosine (5caC).^[8, 9]^ TET-mediated oxidation of 5mC initiates DNA demethylation, which leads to the removal of repressive methylation marks and gene reactivation.

The mammalian TET family consists of three proteins (TET1, TET2 and TET3) that share significant sequence homology. All three TET proteins split molecular oxygen with the help of non-heme iron (Fe (II)): one of the oxygens is inserted into the DNA substrate, while the second one is used to oxidize α-ketoglutarate.^[8, 9]^ However, TET proteins differ in the structure of their DNA binding domain and may have different functions in epigenetic regulation. TET1 and TET3 contain a CXXC domain that aids in their binding to CpG rich sequences. TET2 lacks the CXXC domain. Instead, it partners with IDAX protein, an independent CXXC domain containing protein which is originally encoded by the *TET2* gene and separated through a chromosomal inversion.^[10, 11]^ Studies suggest that the CXXC domain may regulate the genomic distribution of TET proteins. For example, TET1 protein preferentially binds to CpG enriched promoters of gene in mouse embryonic stem cells (mESCs).^[12]^ Similarly, TET3 has slightly higher binding preference for CpG rich gene promoter sequences.^[11]^ In contrast, TET2 has a higher binding preference for gene bodies and enhancer regions rather than gene promoters.^[13]^

The exact functions of TET proteins in cancer are incompletely understood. Expression levels of all three *TET* genes are sharply decreased in many human cancers including lymphoma, liver, breast, pancreas, prostate, and lung neoplasms, with the accompanying reduction in global 5hmC levels in tumors as compared with surrounding tissues.^[14]^ It has been hypothesized that TET proteins counteract sporadic *de novo* methylation of CpG sites, which would otherwise inactivate important tumor suppressor and DNA repair genes.^[15]^ This suggests a possible role of TET proteins in preventing tumorigenesis. Specifically, *TET2* appears to act as a tumor suppressor that is involved in the control of balancing survival, growth, and differentiation in normal hematopoiesis.^[16]^ Indeed, acute deletion of *TET3* in *TET2*-deficient mice has been reported to cause myeloid leukemia.^[17]^ In contrast, 5hmC levels are increased in diffuse intrinsic pontine glioblastoma,^[18]^ and *TET1* is significantly upregulated in MLL-rearranged leukemia, leading to global increase of 5hmC levels.^[19]^ These apparently contradictory results limit our understanding of the functional roles of TET proteins and cytosine hydroxymethylation in cancer.

Lung cancer is among the deadliest cancer types, with an average 5-year survival rate of only17.7%.^[20]^ Lung cancer is characterized by both genetic mutations and epigenetic deregulation. Epigenetic deregulation including DNA methylation, histone methylation and non-coding RNA expression contribute to lung cancer.^[21]^ Specifically, upregulation of DNA methyl transferases (DNMT) is frequently observed in lung cancer and is independently associated with poor prognosis.^[22, 23]^ The epigenetic roles of TET proteins in lung cancer have not been fully elucidated. TET2 is altered in 4.04% of non-small cell lung carcinomas, with TET2 mutations present in 3.54% of all non-small cell lung carcinoma patients.^[24]^ Recent studies based on cell line models proposed a mechanism where *TET1* gene is downregulated via promoter methylation following exposure to environmental chemicals.^[25, 26]^ On the contrary, a separate study reported that *TET1* is upregulated (2-45-fold) and acts as an oncogene in lung tumors characterized by p53 loss of function.^[27]^ Therefore, a precise role of *TET1* in human lung cancer remains to be elucidated.

In the present study, we employed CRISPR-Cas-9 gene editing and molecular cloning methodologies to modulate the levels of *TET1* gene expression in cell culture and 3D spheroid models of lung cancer. Our results indicate that *TET1* acts as a tumor suppressor gene in human lung adenocarcinoma. *TET1* overexpression decreases tumor cell proliferation and halters tumor growth, while *TET1* deficiency leads to increased lung tumor growth. Our results reveal the antitumor function of *TET1* in lung cancer and show that *TET1* induces innate immunity through activation of the TNF signaling pathway.

## Results

### *TET1* expression in lung cancer

5-hmC loss is an epigenetic signature of many cancers, with diagnostic and prognostic implications. Reduced expression of the three *TET* genes has been associated with a decrease in the 5-hmC levels in various cancers, suggesting a plausible mechanism to explain 5-hmC loss in cancer cells. In addition, it has shown that reduced levels of 5-hmC are linked to tumorigenesis in genetically engineered mouse models of various types of human malignancies.^[14]^ Alrehaili et al. previously reported that the expression levels of all three *TET* genes were significantly downregulated in NSCLC tissues in comparison to their matched normal tissues. These authors reported that *TET1*-mRNA levels were 38.48 ±16.38 in non-small cell lung cancer NSCLC vs. 80.65 ± 11.25 in normal tissues (t = 21.33, p < 0.0001).^[28]^

Previous meta-analysis of published studies reveal that high expression levels of the *TET1* gene in solid tumors including lung adenocarcinoma are associated with better overall survival (HR = 0.778, 95% CI = 0.639–0.946, *P* = .012).^[29]^ We were interested in seeing whether there was any evidence for *TET1* as a tumor suppressor within lung cancer. To do this, we utilized the open-source Kaplan-Meier plotter devised by Gyorffy.^[30]^ We selected the lung cancer cohorts originally taken from the NCBI Gene Expression Omnibus and Genomic Data Commons Data Portal and allowed for adenocarcinomas in the histology setting and left all other controls in their default setting. The program then generated the Kaplan-Meier plot seen in Figure 1 **(Supplementary Figure S1)**. While the p-value suggested that there wasn’t a statistically significant difference between high and low expression of *TET1* (P = 0.06), the traces in the plot indicate a slightly higher rate of survival for high *TET1* expression patients over low *TET1* expression patients, suggesting that *TET1* may act as a tumor suppressor in lung cancer.

**Figure 1.**
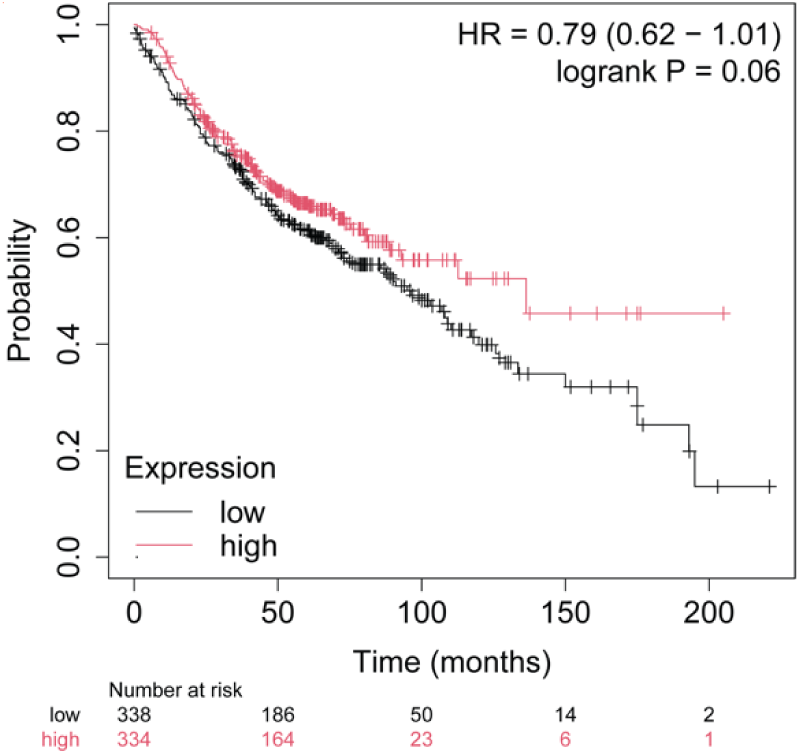
Lung adenocarcinoma for both male and female patients’ survival as a function of *TET1* gene expression.

### TET1 blocks lung adenocarcinoma cancer cell growth in vitro

To directly assess the functional role of *TET1* in lung cancer, we overexpressed TET1 protein in human lung adenocarcinoma cells lines (H441 and H1975). The overexpression efficiency was confirmed by Liquid chromatography-mass spectrometry (LC-MS) assay and western blotting (**Figure 2A and Supplementary Figure S2a-d**). Isotope dilution HPLC-ESI-MS/MS assays revealed that the global levels of 5-hmC in genomic DNA were elevated 3-fold upon TET1 overexpression (**Figure 2B**). Furthermore, TET1 overexpression led to decreased levels of cell proliferation (**Figures 2C, Supplementary Figure S3a**) and reduced colony formation in H441 and H1975 cells (**Figures 2D and S3b**). Trans-well cell migration assays in H441 cells revealed a significant decrease in cell migration in cells overexpressing *TET1*, suggesting that it decreases cell migration and invasion (**Figure 2E**). Next, we employed three-dimensional *in vitro* tumor spheroid models to evaluate whether *TET1* has a role in tumor growth in lung cancer. This experiment revealed that 3D spheroids overexpressing *TET1* grew at a significantly slower rate as compared to control tumors with basal *TET1* expression levels (**Figure 2F and Supplementary Figure S4**). This result is consistent with strong inhibition of lung tumor cell growth observed in 2D cultures (**Figure 2c**). Overall, these results indicate that *TET1* acts as a tumor suppressor gene in lung adenocarcinoma.

**Figure 2.**
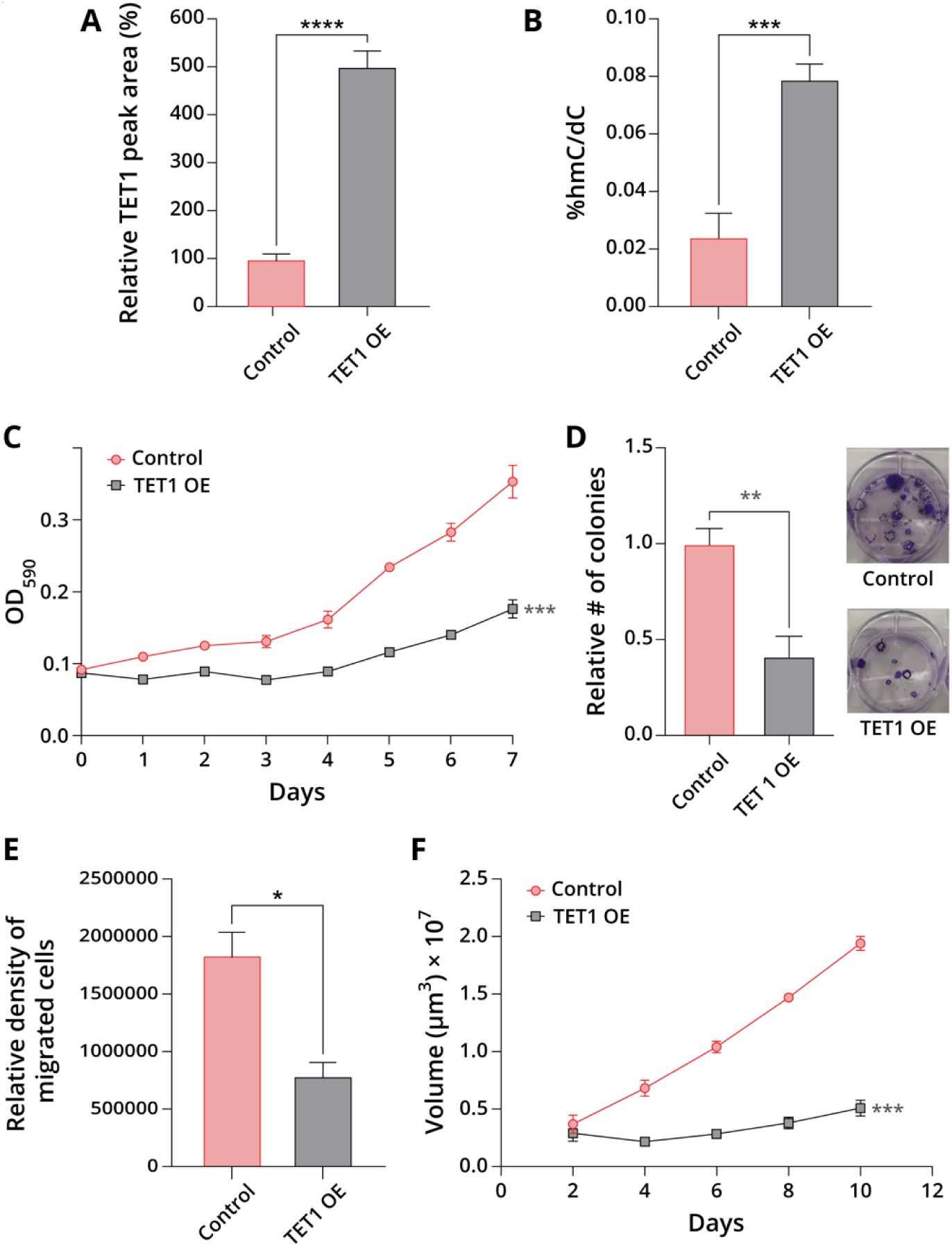
*TET1* overexpression in H441 cells leads to tumor suppressive phenotype. **a,** LC-MS assay to show the relative levels of TET1 protein in control and *TET1* overexpressing cells. **b,** Global levels of 5hmC in H441 cells upon *TET1* OE. Data are expressed as percent of dC and represents mean values ± SD of N = 3. *p < 0.05, **p < 0.01, ***p < 0.001. **c,** Cell proliferation of control and *TET1* overexpressing H441 cells**. d,** Colony formation assays in control and *TET1* overexpressing H441 cells. **e,** Trans-well cell migration assays in control and *TET1* overexpressing H441 cells. **f,** 3D spheroid tumorigenic studies in control and *TET1* overexpressing H441 cells.

### TET1 OE is associated with high levels of immune and antiviral response markers

To help unravel the molecular mechanisms responsible for *TET1* tumor suppressing effects in lung adenocarcinoma, we conducted RNA-seq analyses in *TET1* overexpressing H441 cells. These analyses revealed that *TET1* alters the expression of over 5300 genes, with about 70% genes showing increased levels of expression (**Supplementary Figure S7a**). Among these, MX1 (MX dynamin like GTPase 1) has antiviral activity against a wide range of RNA viruses and some DNA viruses.^[31]^ OAS2 (2’-5’-oligoadenylate synthetase 2) is involved in the innate immune response to viral infection.^[32]^ Additionally, OAS2 may also play a role in other cellular processes including apoptosis, cell growth, differentiation, and gene regulation.^[33, 34]^ RASD2 (radical S-adenosyl methionine domain containing 2) is an interferon-inducible gene that plays a role in cellular antiviral response and innate immune signaling.^[35]^ Another important gene activated upon *TET1* overexpression is XAF1, a tumor suppressor gene which mediates TNFα-induced apoptosis.^[36, 37]^ (**See Supplementary Figure S7a**).

Gene ontology (GO) analysis of genes showing increased expression levels in cells overexpressing *TET1* revealed a total of 36 GO terms including TNF signaling pathway, NF-kB signaling pathway, IL-17 signaling pathway, apoptosis, and necroptosis pathways (**Figure 3A and Supplementary Figure S7c**). Gene interactome analysis for IL-17 signaling pathway revealed over 30 genes associated with each other, with TNF and NFkB1 at the center of the cluster and interacting with most of the other genes (**Figure 3B**). TNF is a multifunctional cytokine that plays vital roles in various cellular processes such as immune response, inflammation, proliferation, apoptosis etc.^[38]^ NFkB1 is one of the five members of the NF-kB family which regulates various processes such as transcription, inflammation and immune responses.^[39, 40]^ Gene interactome analysis of the apoptosis pathway showed CASP8 gene associating with most of the other genes in the cluster (**Figure 3C**). Activation of CASP8 is known to have a central role in cell apoptosis.^[41]^ CASP7 has also been enriched in the cluster which is known to facilitate the execution of apoptosis through downregulation of the 26S proteosome.^[42]^ Further, we analyzed the top 10 genes with the largest logFC values in the gene clusters associated with their GO terms. We chose the TNF signaling pathway, NF-kB signaling pathway, IL-17 signaling pathway, necroptosis, and apoptosis. These analyses revealed that the top 10 genes in the KEGG reactome pathways are immune marker genes such as CXCL10, CCL5, INFB1, CCL20, TNF, LBT, and others (**Figures 3D-E and supplementary Figure S9a-c**). Overall, these results indicate that *TET1* overexpression in H441 lung cells activates an immune response via the TNF-alpha signaling pathway.

**Figure 3.**
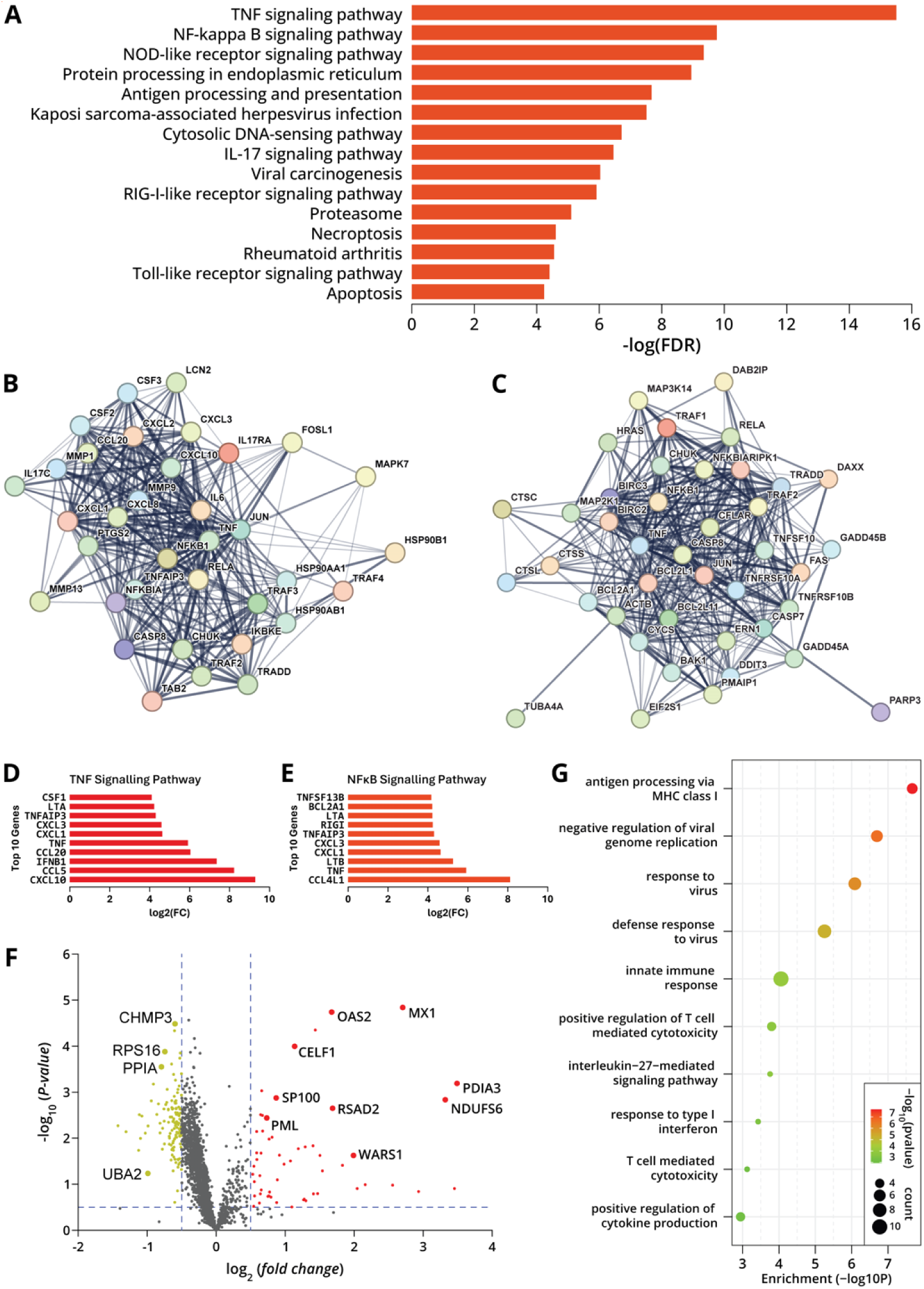
Transcriptomic and proteomic studies of *TET1* overexpressing H441 cells. **a,** Top 15 KEEG biological pathways of significantly enriched upregulated genes. **b-c,** Gene clustered associated with *TET1* OE in IL-17 and apoptosis pathways. **d-e,** Top 10 enriched genes in TNF and NFkB signaling pathways. **f,** Volcano plot of quantitative analysis of proteins identified upon *TET1* OE in H441 cells. **g,** Dot-plot diagram of upregulated proteins showing top 10 biological processes upon *TET1* OE in H441 cells.

Next, we employed mass spectrometry-based global proteomics to investigate the changes in protein levels in H441 cells upon *TET1* overexpression. TMT6-plex based proteomics identified over 2300 protein groups changing in abundance upon *TET1* overexpression (**Figure 3F**). Among these, 60 proteins were significantly increased in abundance, and 167 proteins decreased in abundance (**Figure 3F**). RPS16, PPIA, CHMP3, and UBA2 are among the most significantly decrease in abundance. While MX1, OAS2, CELF1, PDIA3, NDUFS6, RSAD2, WARS1, and SP100 were among the most significantly increased in abundance in *TET1* OE cells (**see volcano plot in Figure 3F**). This is fully consistent with our RNA-seq results discussed above. GO enrichment analysis of the proteins increased in abundance in *TET1* OE cells revealed a number of important biological processes including defense response to virus, innate immune response, and the NF-kappa B signaling pathway (**Supplementary Figure S10a**). Similar pathways were found to be upregulated in RNA-seq analysis of *TET1* OE cells (**Figure 3A-C**). The dot-plot diagram for the top 10 biological processes shows that the majority of upregulated proteins in *TET1* overexpressing H441 cells are involved in antiviral and immune responses (**Figure 3G**). Collectively, our multi-omics results indicate that *TET1* OE induces immune response and antiviral response pathways in H441 lung adenocarcinoma cells.

### CRISPR-cas9 to generate *TET1* KO in H441 lung cells

The above results indicate that *TET1* OE blocks pulmonary adenocarcinoma cell proliferation and induces immune responses through activation of TNF signaling pathway. Next, we investigated the effects of *TET1* KO in lung adenocarcinoma (H441) lung cancer cells. This cell line was selected because other *TET1* expressing lung adenocarcinoma cell lines (e.g. H2023, H358, H1435, H1993 etc.) have excessive multiploidy and are not amenable to CRISPR.^[27]^ We have created four *TET1* KO clones (clone 2, 4, 23, 24) in H441 cells with CRISPR/cas9 targeting exon4 of the *TET1* gene (**Figure 4A**). Genotyping studies showed that *TET1* KO using both 5’ and 3’ gRNA results in 1873 bp deletion and generation of 427 bp with a premature stop codon (**Figure 4A, Supplementary Figure S5**). Western blotting analysis showed successful KO of TET1 (**Figure 4B and Supplementary Figure S6a**). To investigate the outcome of *TET1* knockout in H441 cells, we carried out isotope dilution HPLC-ESI-MS/MS analysis of the global levels of 5mC and 5hmC in *TET1* KO and H441-wt cells. Global 5-methylcytosine (5mC) levels in genomic DNA isolated from *TET1* KO clones were similar to that in wild type cells (**Figure 4C**). In contrast, 5-hydroxymethylcytosine (5hmC) levels were significantly lower in DNA isolated from clones 23 and 24 (**Figure 4D**), consistent with reduced levels of TET activity. To determine whether *TET1* KO leads to changes in the expression levels of TET2 in an attempt to compensate for TET1 loss, we also conducted western blotting for the TET2 protein in *TET1* KO clone. TET2 western blotting revealed that there was no significant change in the expression levels of TET2 in *TET1* KO clone compared to the H441-WT cells (**Supplementary Figure S6b)**. Based on these results, *TET1* KO clones 23 and 24 were selected for functional analyses.

**Figure 4.**
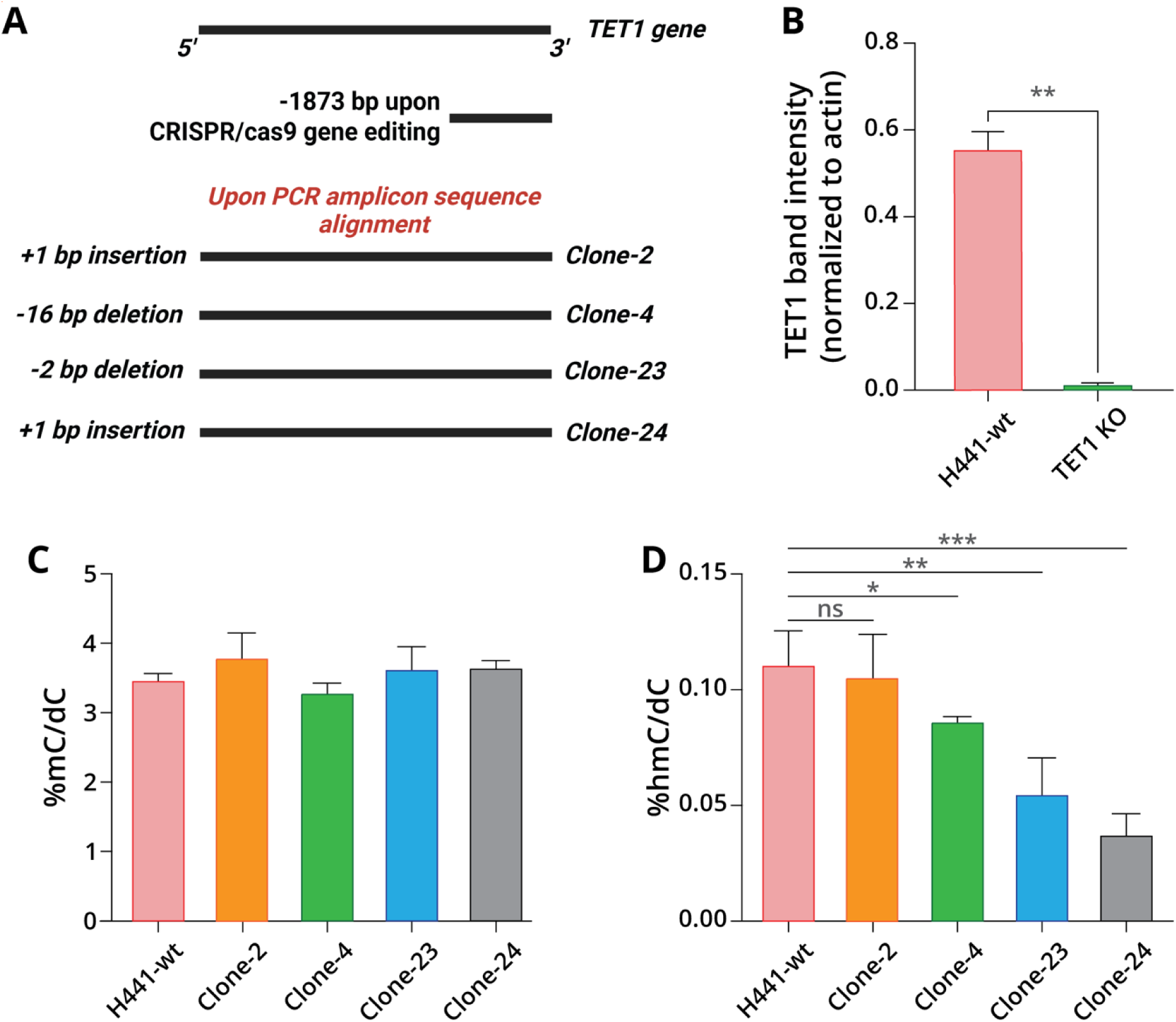
CRISPR/cas9 gene editing of *TET1* gene in H441 cells and characterization of KO clones. **A,** Pictorial representation of four *TET1* KO clone variants. **B,** TET1 western blot characterization of *TET1* KO. **C,** Global levels of 5-methylcytosine in *TET1* KO cells through LC-MS analysis. Data represents mean values ± standard deviation (SD); N = 3. **D,** Global levels of 5-hydroxymethylcytosine in *TET1* KO cells. Data represents mean values ± standard deviation (SD); N = 3.

### *TET1* KO in H441 LUAD cells promotes cell proliferation

To evaluate the phenotypic effects of *TET1* KO in lung adenocarcinoma cells, we carried out proliferation assays and observed that clones 23 and 24 had increased rates of cell proliferation as compared to H441-wt cells, consistent with *TET1* gene has a role in tumor suppressor (**Figure 5A**). In contrast, cell proliferation rates of clones 2 and 4 were similar to wild type cells (**Figure 5A**). To further validate this data, we conducted colony formation assays and demonstrated that clones 23 and 24, but not clone 2, had higher number of colonies than H441-wt cells (**Figure 5B**). Trans-well migration assay conducted with clone 24 showed significant increase in cell migration in *TET1* KO cells as compared to controls (**Figure 5C**). Therefore, clone 24 was selected for further studies.

**Figure 5.**
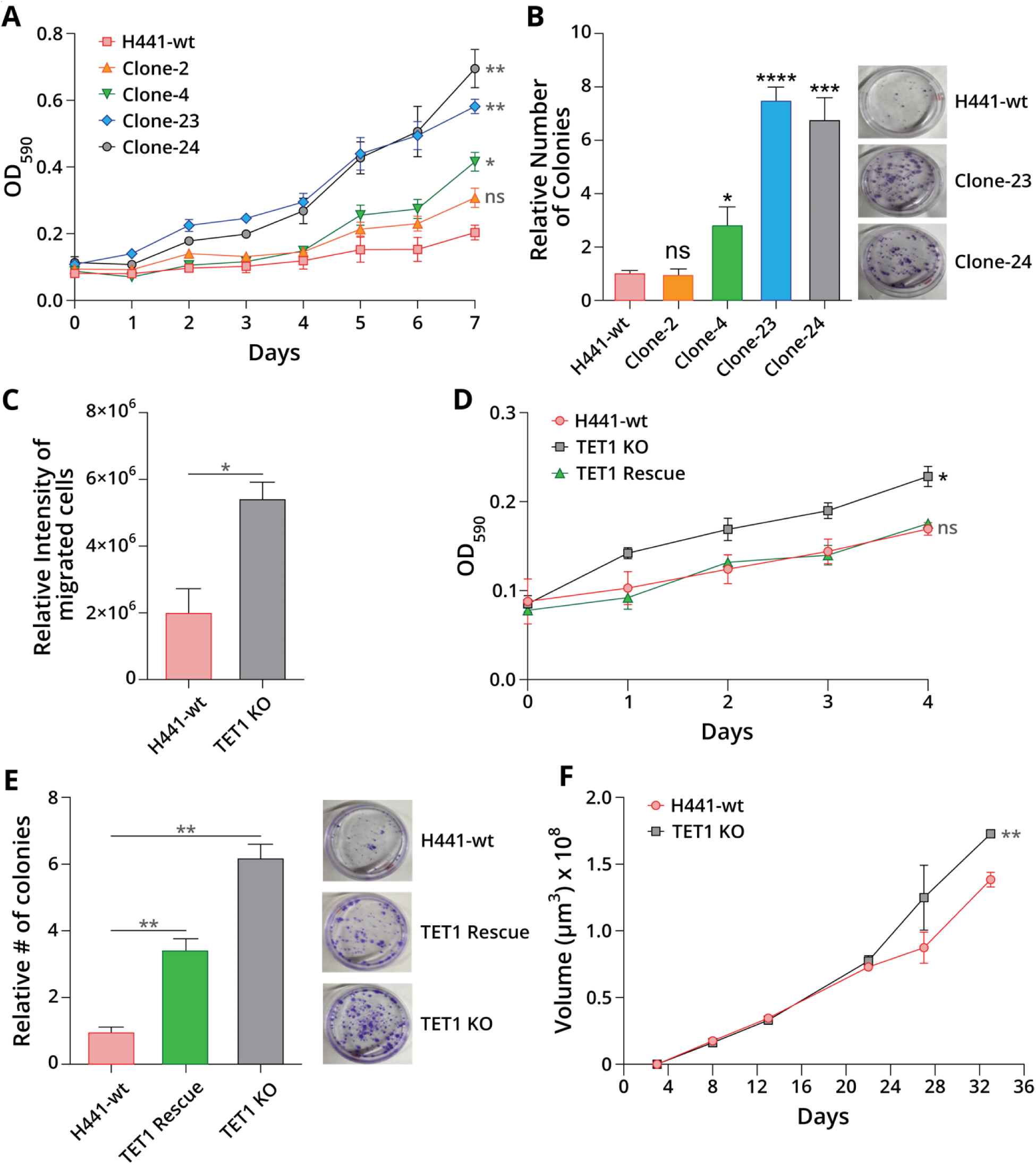
*TET1* knockout in lung adenocarcinoma (H441) cells leads to more aggressive tumor phenotype. **A.** Cell proliferation of *TET1* KO cells as compared to wild type H441 cells. **B,** Relative numbers of colonies in *TET1* KO cells as compared to wild type H441 cells. **C,** Trans-well cell migration assay in *TET1* KO cells. **D,** Cell proliferation for *TET1* re-expressing cells in *TET1* KO cells. **E,** Relative number of colonies for *TET1* re-expression in *TET1* KO cells. **F,** in-vitro 3D spheroid tumorigenic studies in *TET1* KO cells.

To confirm whether the observed phenotypes were a result of *TET1* KO, we carried out TET1 rescue experiments in *TET1* KO cells. To this end, we transiently expressed the *TET1* gene in *TET1* KO H441 lung cells. Cell proliferation data revealed that re-expression of TET1 protein in *TET1* KO cells induced a decrease in cell growth and generated a rescued a cell phenotype similar to H441-wt (**Figure 5D**). Similarly, we carried out a colony formation assay and observed a significant decrease in colony formation in rescued cells as compared to *TET1* KO cells (**Figure 4E**). Although the number of colonies in *TET1* re-expressing cells are higher than H441-wt cells, it is probably because the transfected cells have been monitored over 3 weeks for colony formation. And moreover, after a certain time cells will slowly come back to their original phenotype in transient transfection. Thus, *TET1* re-expression in *TET1* KO cells induced a recovery of the cell phenotype, confirming that the increase in cell proliferation in *TET1* KO cells is attributed to *TET1* deficiency. Further, we investigated 3D spheroid tumorigenesis of *TET1* KO cells in vitro. 3D spheroid assays revealed that the growth of 3D spheroids of *TET1* deficient cells was significantly faster than for H441-wt cells after 3 weeks of growth (**Figure 4F**). Taken together, these results are consistent with *TET1* acting as a tumor suppressor gene in H441 LUAD cells.

### *TET1* KO induces translation and biosynthetic pathways

To identify potential mechanisms involved in more aggressive cancer phenotype in lung adenocarcinoma cells lacking the *TET1* gene, global transcriptomics and proteomics experiments were conducted. RNA-seq data revealed that there were over 1500 differentially expressed genes (DEG), of which about 40% genes were upregulated in *TET1* KO cells (**Supplementary Figure S7b**). EIF3H, SLC9A9, HLA-DRA, and CD74 were among the most significantly enriched genes, while TUSC3, PRSS21, MT1E, and DDX3Y were the most significantly downregulated genes in *TET1* KO cells (**Supplementary Figure S7b**). Literature reports suggest that EIF3H is associated with protein biosynthesis and initiates translation of a subset of mRNA which involves in cell proliferation, differentiation, and cell cycle.^[43–45]^ This is consistent with accelerated cell growth and proliferation in *TET1* KO cells. Among downregulated genes, TUSC3 functions as a tumor suppressor gene in many cancers. For example, downregulation of TUSC3 has been shown to promote hepatocellular carcinoma through LIPC/AKT axis.^[46, 47]^

GO enrichment analysis carried out on genes upregulated in *TET1* KO cells revealed a total of 38 GO terms including eukaryotic translation, peptide chain elongation, metabolism of amino acids derivatives and ribosomal assembly (**Figure 6A, Supplementary Figure S7d**). Next, we conducted gene interactome analysis on upregulated genes and observed various RPL and RPS genes co-expressed and formed a cluster at the center of the interactome (**Supplementary Figure S8**). Overexpression of these genes in *TET1* KO cells leads to translation and various biosynthetic processes which have been observed in GO terms. Further, we analyzed the top 10 genes with the largest logFC values associated with their GO terms. We chose peptide chain elongation, cellular response to stimuli, signaling by ROBO receptors, metabolism of amino acids and derivatives, and disease pathways (**Figures 6B-C and Supplementary Figure S9d-f**). In the peptide chain elongation, all the top 10 genes contribute to protein biosynthesis. For pathways involved in metabolism of amino acids and derivatives, we observed overexpression of *NMMT, CPS1, IDO1, RPS26*, and other genes that are involved in metabolism of amino acids such as tryptophan, and sarcosine.^[48]^ Sarcosine has been prevuously identified in metabolomics studies as a biomarker of prostate cancer progression.^[49]^

**Figure 6.**
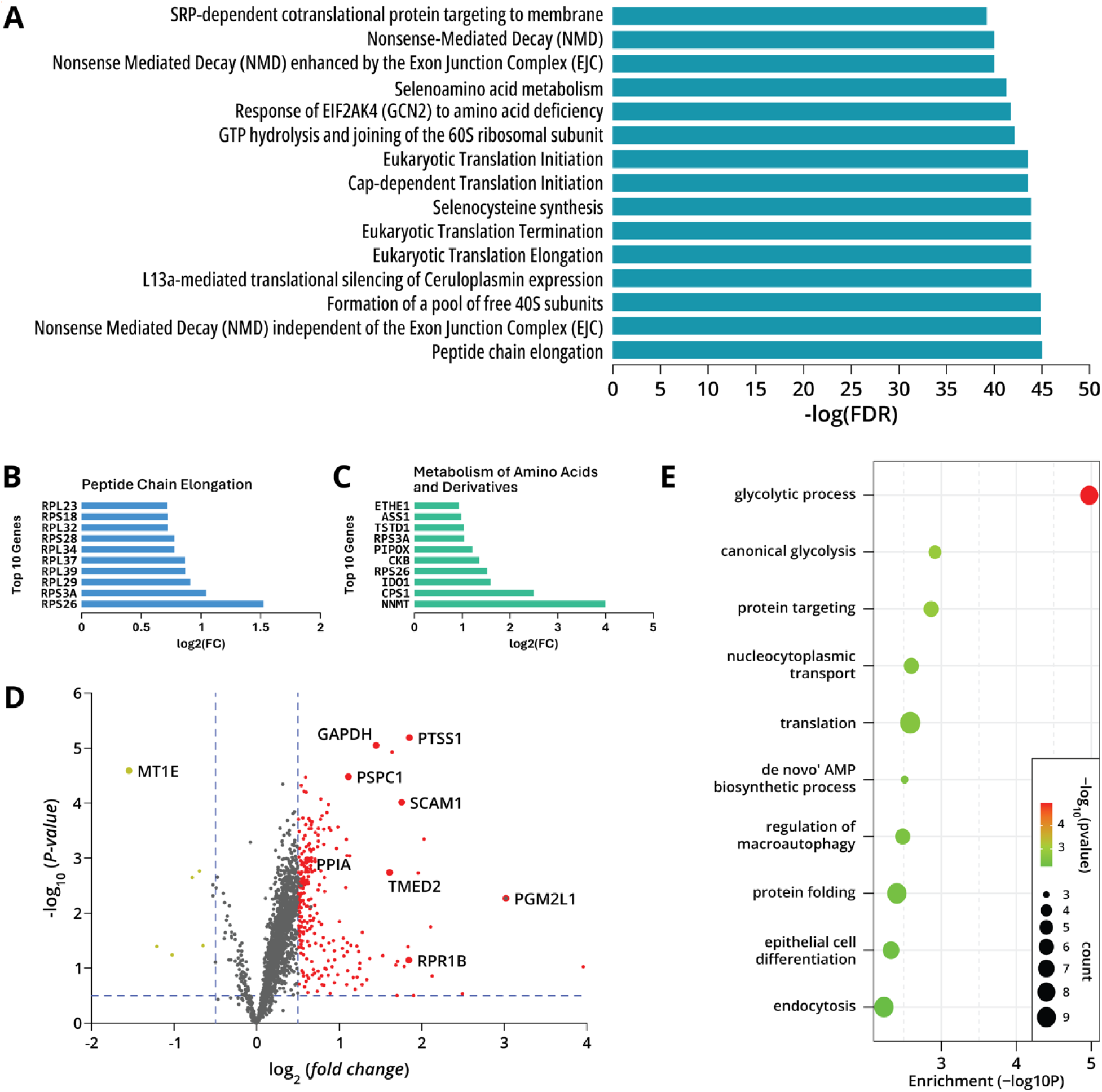
Transcriptomic and proteomic changes in *TET1* KO H441 cells. **a,** KEEG biological pathways of significantly enriched upregulated genes. **b,** Top 10 enriched genes in peptide chain elongation pathway. **c,** Top 10 enriched genes in metabolism of amino acids and derivatives pathway. **d,** Volcano plot of quantitative analysis of proteins identified upon *TET1* KO in H441 cells. **e,** Dot-plot diagram of upregulated proteins showing top 10 biological processes upon *TET1* KO in H441 cells.

Next, we investigated global protein changes in *TET1* deficient H441 cells. Mass spectrometry based quantitative proteomics identified over 2100 protein groups. Among these the majority were significantly increased in abundance and only 6 proteins were downregulated. GAPDH, PTSS1, SCAM1, TMED2, PGM2L1, PSPC1, and RPR1B were some of the most significantly upregulated proteins in *TET1* deficient cells (**Figure 6D**). GO enrichment analysis from the upregulated proteins revealed various important biological processes such as glycolysis, translation, and *de novo* AMP biosynthesis (**Supplementary Figure S10b**). Similar biosynthetic and translational pathways have also been observed in RNA-seq data of *TET1* deficient cells. These biosynthetic and glycolysis pathways are involved in the production of biomolecules necessary for cell growth, which is consistent with increased proliferation of *TET1* KO cells. A dot-plot diagram (**Figure 6E**) shows that most of the upregulated proteins in *TET1* KO cells are involved in translation, glycolysis, protein targeting, and various biosynthetic processes. Taken together, transcriptomics and proteomics results indicate that *TET1* deficiency in lung adenocarcinoma (H441) cells induces translation, glycolysis, and various metabolic and biosynthetic pathways which may produce the necessary energy and biomolecules required for cell proliferation and tumor growth.

## Discussion

In this study, *TET1* is demonstrated as a tumor suppressive gene in human lung adenocarcinoma cells. We found that *TET1* overexpression in lung adenocarcinoma (H441 and H1975) cells blocks cell proliferation, colony formation, and cell migration **(Figure 2**). *In vitro* 3D spheroid assay revealed that *TET1* OE blocks the growth of tumor spheroids. In contrast, *TET1* deficiency in H441 cells promoted cell proliferation, colony formation, and cell migration (**Figure 5**). *TET1* deficiency increases the growth of 3D lung adenocarcinoma tumor spheroids. These results are consistent with previous reports that *TET1* knockout in transgenic mice induces tumor growth.^[50]^

Transcriptomics analyses of H441 cells overexpressing *TET1* revealed upregulation of genes associated with immunity and antiviral responses. Among the most significant upregulated genes, *MX1, RSAD2, OAS2, XAF1* mainly contribute to innate immune responses, apoptosis, and antiviral activity against various viruses.^[31–35, 37]^ GO enrichment analysis revealed that the upregulated genes primarily activate TNF signaling pathway, NF-kB, apoptosis etc. For example, NF-kB family members are known to control the transcription of cytokines, antimicrobial effectors as well as genes that regulate proliferation, differentiation and regulate various aspects of innate immune responses.^[51]^ Gene interactome analyses reveal that co-expression of various immune markers with the TNF and NF-kB1 clustered at the center. The top 10 genes in various upregulated pathways revealed that most of the genes are immune marker genes. Quantitative proteomics analyses on *TET1* OE H441 cells revealed similar sets of upregulated proteins whose encoding genes were observed in RNA seq. data. GO enrichment analysis on upregulated proteins revealed the activation of innate immunity, NF-kappa B signaling pathway, and various antiviral responses pathways. Overall, transcriptomics and proteomics data suggest that *TET1* OE in H441 lung cells reduces tumorigenesis through the activation of antitumor immunity.

Multi-omics studies of lung adenocarcinoma cells lacking the *TET1* gene revealed activation of translational and biosynthetic pathways (**Figure 6**). Among top upregulated genes, *EIF3H* is associated with protein biosynthesis and initiates translation of a subset of mRNA which involves in cell proliferation, differentiation, and cell cycle.^[43, 44]^ Literature reports suggest that elevated levels of *EIF3H* have been found in lung adenocarcinoma.^[52]^ Another upregulated gene, *SLC9A9,* is a membrane protein that regulates the luminal pH of the recycling endosome and also reported oncogenic in various cancers such as esophageal squamous carcinoma, glioblastoma and colorectal cancer.^[53]^ However, to our knowledge there are no previous reports of the oncogenic role of *SLC9A9* in lung cancer. On the other hand, among top downregulated genes *TUSC3* has been reported to promote HCC, cervical squamous cell carcinoma.^[46, 54]^ However, it has been reported that downregulation of *TUSC3* in NSCLC (H1299, A549) decreases cell proliferation, cell migration, invasion via upregulation of claudin-1.^[55]^ Overall, our data reveals that *TET1* KO leads to upregulation of various genes that may enrich various translational and biosynthetic pathways which may contribute to increase in cell proliferation, colony formation, and cell migration in H441 cells. Proteomics data for *TET1* KO in H441 cells mainly enriched glycolysis, translation, and various biosynthetic pathways in upregulated proteins. Similar sets of pathways have also been observed in RNA seq. data of *TET1* KO cells.

One of the major pathways observed from proteomics data in *TET1* KO cells is glycolysis which is an important metabolic property of tumor cells that provide rapid energy, an essential precursor for a variety of other metabolic pathways and raw materials for the synthesis of multiple biomolecules.^[56]^ Moreover, glycolysis has been shown to promote tumor invasion and migration in pancreatic ductal adenocarcinoma.^[57]^ In the current study, proteomics data revealed that *TET1* KO enriched glycolysis, translation which may increase tumorigenesis in H441 lung cells.

In conclusion, our data reveal that *TET1* functions as a tumor suppressor gene in human H441 and H1975 lung cancer cells. We have conducted a systematic functional analysis through the modulation of *TET1* gene in H441 and H1975 to show its tumor suppressor role. Transcriptomics and proteomics analysis identified TNF mediated activation of immunity in *TET1* overexpressing H441 cells. From the link between *TET1* overexpression and immunity and given the importance of immune infiltration in clinical outcomes, it is worth rethinking how *TET1* expression relates to cancer. In the future one may explore the use of epigenetic drugs modulating antitumor immune responses in order to dissect the epigenetic mechanisms underlying the crosstalk between the immune system and cancer immunotherapy.

## MATERIALS AND METHODS

### Cell culture

All cells were acquired from ATCC (www.atcc.org). H441 and H1975 cells were cultured in RPMI media (catalog number: 22400-089) supplemented with 10% FBS and 1% penicillin-streptomycin at 37 °C in a humidified 5% CO_2_ and sub-cultured for up to passage number 7.

### Transfection

H441 and H1975 cells were transfected using lipofectamine 3000 transfection reagent (Invitrogen, L3000-008) according to the manufacturer’s protocol. After 40 h of transfection, cells were isolated, counted and seeded for cell proliferation, colony formation, and migration assays.

### MTT cell proliferation assay

4000 cells were seeded in each well in 100 uL of RPMI media supplemented with 10% FBS and 1% penicillin-streptomycin of a 96 well plate. There are three biological replicates and three technical replicates for each biological replicate. After cells were attached, added 10 uL of MTT reagent (cat. 11465007001) in each well and kept for incubation at 37 °C for 4 h. After 4 h, MTT solvent (150 uL) was added to each well and incubated overnight at 37 °C. Next day morning, measured the absorbance at 590 nm in a plate reader.

### Colony formation assay

2000 cells for each replicate were seeded in a 6 cm dish in 5 mL of RPMI1640 media (two technical replicates for each biological replicate) or 1000 cells were seeded per well in a 6 well plate in 2 mL of media and kept for incubation at 37 °C. Media was replaced in every 3-4 days. After around 3 weeks (when visible colonies were observed), the media was aspirated off and the cells washed 1X with PBS and fixed with 0.6% crystal violet in 6% glutaraldehyde for 30 minutes at room temperature. After 30 minutes, the fixing media was aspirated off and the cells were washed with water and let the plate dry completely. The colonies were then counted manually.

### Trans-well cell migration assay

Cell suspensions were prepared in serum free media. 50,000 cells in 300 uL serum free media were seeded onto the apical side of the cell culture inserts (cat. PTEP24H48) in the 24 well plate. 750 uL media supplemented with 10% FBS was added to the basolateral side of each insert in the wells. An insert without cells was included as a blank control and an insert with cells and media without FBS in the basolateral side of the insert was included as a negative control. After 72 h, media was removed and the cells washed 2X with PBS and the cells fixed with 4% formaldehyde for 5 mins at room temperature. After fixation, the cells were washed 2X with PBS and were then permeabilized in 100% Methanol for 20 minutes at room temperature. The methanol was then removed, and the cells were washed twice with PBS, after which the cells were stained with 0.5% crystal violet for 15 minutes at room temperature. The stained cells were washed with PBS and allowed to dry, after which cells on the apical side were removed carefully using cotton swabs. Cells at basolateral side of the inserts were considered to be migrated cells.

### TET1 Western blotting

100 uL lysis buffer (7M urae, 2M thiourea, 10 mM TCEP, 40 mM 2-chloroacetamide, 40 mM Tris base, 20% ACN, protease inhibitor) was added to cell pellet (∼5 million). Cell pellets were gently resuspended via pipette and then vortexed for 2-3 minutes. Cell suspension was then sonicated for 15 seconds 1 sec. OFF and 1 sec. ON with 30% amplitude. Cell lysates were then incubated for 30 minutes at 37 °C. Denatured protein lysates (50 ug) were separated by SDS gel electrophoresis and transferred to nitrocellulose membranes (Invitrogen) for 1 h at 30 V. After blocking in 1% BSA solution for 1 h at room temperature, the membranes were incubated overnight with the primary antibody (Cat No. GTX124207) at 4 °C. This was followed by incubation with their respective IR-dye secondary antibodies and the signals were visualized using LiCor imager.

### 3D spheroid assay

The 3D Petri Dishes were prepared according to the manufacturer’s protocol with 2% ultra-pure agarose in 0.9% NaCl from a 24-35 series micro mold purchased from microtissues. Next, 3D Petri Dishes were transferred to 12 well plates. To equilibrate the 3D Petri Dishes, 2.5 mL cell culture media in each well was added and incubated for 15 minutes. The media was removed and replaced with fresh media and incubated for another 15 minutes. Next, all the media from the well and from the cell seeding chamber of the 3D Petri Dishes was carefully removed. 35,000 cell/75 uL of media were seeded dropwise into the cell seeding chamber. Cells were allowed to settle into the features of the 3D Petri Dishes and then additional media (2.5 mL) were added to the outside of the 3D Petri Dish. Next, place the tissue culture plate in the cell culture incubator at 37 °C. Then over the time spheroid growth was recorded in a Nikon Ti-E Deconvolution Microscope System and media was exchanged every 3-4 days.

### DNA and RNA Extraction

DNA and RNA were extracted from cells by Qiagen All prep DNA/RNA mini kit according to the manufacturer’s instructions. DNA concentration was calculated using Qubit 4 fluorometer. RNA was stored immediately at −80 °C after extraction. DNA was stored at −20 °C.

### RNA seq analysis of H441 OE and knock out cells

After the extraction and quantification of RNA, RNA integrity was confirmed using Agilent Bioanalyzer (Agilent, Santa Clara, CA, USA). RNA was reverse transcribed into cDNA and Illumina adapters were added using PCR. Sequencing of samples was performed on the Illumina NovaSeq 6000 sequencing system.

### RNA seq read processing

RNA-Seq. analyses were performed within the Galaxy bioinformatics suite.^[58]^ FASTQ files were processed via Trimmomatic v0.38.1^[59]^ to remove Illumina sequencing primers and low-quality reads. The resulting files were then aligned with *Homo sapiens* GRCh38.87.gtf gene annotation file using HiSAT2 v2.1.0 ^[60]^ to generate BAM files; these BAM files and the gene annotation file were then used by featureCounts v2.0.1 ^[61]^ to generate counts for RNA-Seq reads. The resulting counts files were then fed into edgeR v3.24.1,^[62]^ wherein differential analysis was performed using an adjusted p-value threshold of 0.05 for change and trimmed mean of m-values normalization. The resulting logFC values and FDR values were used to generate volcano plots to illustrate the differential expression of genes.

### RNA seq gene expression quantification and filtering

Genes showing significant increases or decreases in abundance were analyzed using the gprofiler suite to perform Gene Ontology analyses.^[63]^ The genes with the ten largest logFC values in the gene clusters associated with their GO terms were plotted to illustrate the contents of the gene clusters.

### HPLC-ESI-MS/MS Quantitation of 5mC and 5hmC

Quantitation of 5hmC is based on methodology previously described by our group in Seiler et al. with modification.^[64]^ Genomic DNA (1-2 ug) was dried down and spiked with 1 pmol each of 5-Methyl-2’-deoxycytidine-d_3_ and 5-(Hydroxymethyl)-2’-deoxycytidine-d_3_. Nucleotide digestion enzyme mix, and buffer were added per product guidelines on a 20 µL scale. Samples were digested for 60 min at 37 °C on a dry heating block. Following digestion, samples were filtered on nanosep 10K filters. Hydrolysates were transferred to MS vials, dried completely, and resuspended in 100 µL mQ-water for offline HPLC cleanup to enrich 5mC and 5hmC. An Atlantis T3 column (Waters, 4.6 x 150 mm, 3 µm) was used with a flow of 0.9 mL/min with a linear gradient of 5 mM NH_4_CHO_2_ buffer, pH 4.0 (Solvent A) and MeOH (Solvent B). Fractions containing 5mC, and 5hmC were collected and pooled before analysis by isotope dilution HPLC-ESI-MS/MS. Pooled samples were dried completely and reconstituted in 2 mM ammonium formate buffer (NH_4_CHO_2_) with a Dionex Ultimate 3000 UHPLC (Thermo Fisher) interfaced with a Thermo TSQ Quantiva mass spectrometer (Thermo Fisher) flowing 2 mM NH_4_CHO_2_ (Solvent A) and ACN (Solvent B) at 15 µL/min through a Zorbax SB-C18 column (Agilent, 0.5 x 150 mm, 3 µm). Quantitation was done monitoring MS/MS transitions m/z 242.1 [M + H]^+^ → m/z 126.1 [M – deoxyribose + H]^+^ for 5mC, m/z 245.2 [M + H]^+^ → m/z 129.0 [M – deoxyribose + H]^+^ for 5-d_3_-mC, m/z 258.1 [M + H]^+^ → m/z 142.1 [M – deoxyribose + H]^+^ for 5hmC, m/z 261.1 [M + H]^+^ → m/z 145.1 [M – deoxyribose + H]^+^ for 5-d_3_-hmC. All mass spectrometry parameters were optimized through direct infusion of authentic standards with settings as follows: spray voltage of 2700 V, sheath gas of 15 au, source fragmentation voltage of 5 V, and ion transfer tube temperature of 350 °C. Fragmentation was induced with an Argon gas flow of 1 mTorr.

### Proteomics Sample Preparation and LC-MS/MS Analysis

#### Sample preparation

Fresh cell pellets were resuspended in cell lysis buffer (8 M urea pH = 8, 50 mM TEAB) supplemented with 1x complete EDTA-free protease inhibitor cocktail. Cell suspensions were then probe sonicated for 15 seconds 1 sec. OFF, 1 sec. ON with 50% amplitude. Cell lysates were then centrifuged at 1600 g for 10 mins. at 4 °C. The supernatant was then transferred into a new tube and the protein concentration was measured with a Bradford assay. 100 ug of protein per condition was transferred into a centrifuge tube and adjusted the volume to 100 uL with 100 mM TEAB buffer. 5 uL of 200 mM TCEP was added and incubated the samples at 55 °C for 1 h. Next, 5 uL of 375 mM iodoacetamide was added to the samples and incubated for 30 mins. at room temperature. 600 uL of pre-chilled acetonitrile was added and the samples and incubated at −20 °C overnight to precipitate out protein. Next day morning samples were centrifuged at 8000 g at 4 °C for 10 minutes. The supernatant was carefully decanted off and the pellet allowed to dry for 2-3 minutes. The pellet was then resuspended in 100 uL of 50 mM TEAB buffer. 2.5 uL (2.5 ug) of trypsin was added into each sample and the samples digested overnight at 37 °C. Digested peptide concentration was measured by Pierce Quantitative Colorimetric Peptide Assay (product No. 23275) and labeled with TMT 6plex according to the manufacturer’s protocol.

Following TMT labeling, the peptides from each sample were concatenated into a single sample. The pooled sample was then desalted and fractionated using the Pierce High pH reverse phase peptide fractionation kit according to the manufacturer’s instruction with a total of 17 fractions. Each fraction was then dried down on speed vac and reconstituted in 10 µL LCMS grade water containing 0.1% formic acid.

#### LC-MS analysis

The seventeen fractionated peptide samples were analyzed on a Q-Exactive mass spectrometer (Thermo Scientific, Waltham, MA) interfaced with a Dionex Ultimate 3000 UHPLC (Thermo Fisher, Waltham,MA, USA). The UHPLC was run in nanoflow mode with a reverse-phase nanoLC column (0.075 mm × 150 mm) created by hand packing a commercially available fused-silica emitter (New Objective, Woburn, MA) with Luna 5 μm C18 (2) 100 Å media (Phenomenex, Torrance, CA). The samples were eluted at a flow rate of 0.25 uL/min with a gradient of 0.1% FA in water (Solvent A) and 0.1% FA in acetonitrile (Solvent B). Solvent composition was linearly changed from 5% to 33% B over 46 mins, then linearly increased to 90% B over the next 5 mins and maintained at 95% B for another 4 mins. The solvent composition was then returned to initial conditions (5% B) and re-equilibrated for 5 mins. The mass spectrometer was operated in positive mode using a Full MS/dd-MS2 experiment with a FWHM of 15 s. In the full scan mode, resolution was 70,000 with an AGC target of 1 × 10^6^, a maximum IT of 200 ms, and a scan range of 400 to 1600 m/z. Tandem mass spectrometry experiments were conducted at 17,500 resolution, AGC target of 1 × 10^6^, maximum IT of 60 ms, an isolation window of 2.0 m/z, a dynamic exclusion time of 30 s, and a normalized collision energy of 30.

### Proteomics Data Analysis

Raw mass spectrometry files were analyzed using Max quant (version 1.6.5.0)^[65]^ in the TMT6plex quantitation mode. The *Homo sapiens* uniprot proteome database was utilized for proteomics analysis. In all instances, carbamidomethylation at cysteine and TMT6 labeling at peptide N-termini and lysine residues were set as static modifications, while methionine oxidation, protein N-terminus acetylation and phosphorylation at serine, threonine, and tyrosine were set as dynamic modifications. Confidence for peptide identifications was set at an FDR cutoff of 0.01. The resulting PSM reports were used for quantitative analysis using Perseus (Version 1.6.5.0).^[66]^ Significantly abundant proteins were tested using student t-test with an FDR cutoff of 0.05. Dot-plot analysis with significantly enriched proteins were carried out with SR plot.^[67]^

### LC-MS assay for TET1 protein

LC-MS/MS with a DDA method and nano emitter packed with C18 matrix was used to analyze the digested TET1 protein for determination of suitable targets for use in a targeted mass spectrometry method. Liquid chromatography using 0.1% formic acid in water (Solvent A) and 0.1% formic acid in acetonitrile (Solvent B) and gradient of 5 to 35% solvent B by 45 minutes. The resulting file were searched against the human proteome (Uniprot, UP000005640) with Fragpipe (version: 22.0, MSFragger version: 4.1, IonQuant version: 1.10.27, Python version: Python 3.9.13), providing peptide-spectral matches with an FDR of 1%. The peptides unique to TET1 were sorted and filtered based on relative intensities, with associated charge states and retention times. This information was implemented in the targeted parallel reaction monitoring (PRM)-based method, using identical LC parameters, with injections of 500 ng acquired for each sample. Peptide peak areas were integrated within Skyline, with MS2 fragmentation characterized by spectral libraries generated using pepXML files of digested TET1 and verified using Koina deep-learning fragmentation prediction. Protein quantitation was summarized by summation of 3 peptides peak areas for each sample, and statistics performed in GraphPad Prism using unpaired, parametric t-tests

