## Supplementary info for "TET1 Functions as a Tumor Suppressor in Lung Adenocarcinoma Through Epigenetic Remodeling and Immune Modulation"

### TET1 is a tumor suppressor gene in lung adenocarcinoma inducing epigenetic remodeling and antitumor immunity

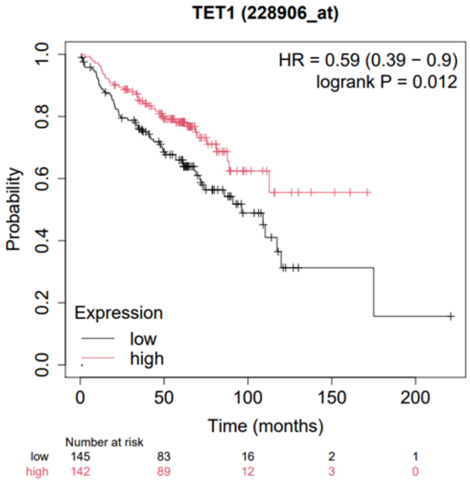


**Figure 1**. Lung adenocarcinoma female patient survival as a function of *TET1* gene expression.

| a) |
|---|
| **TET1 peptide: DQAANEGPEQSSEVNELNQIPSHK, 874.4073+++** |
| 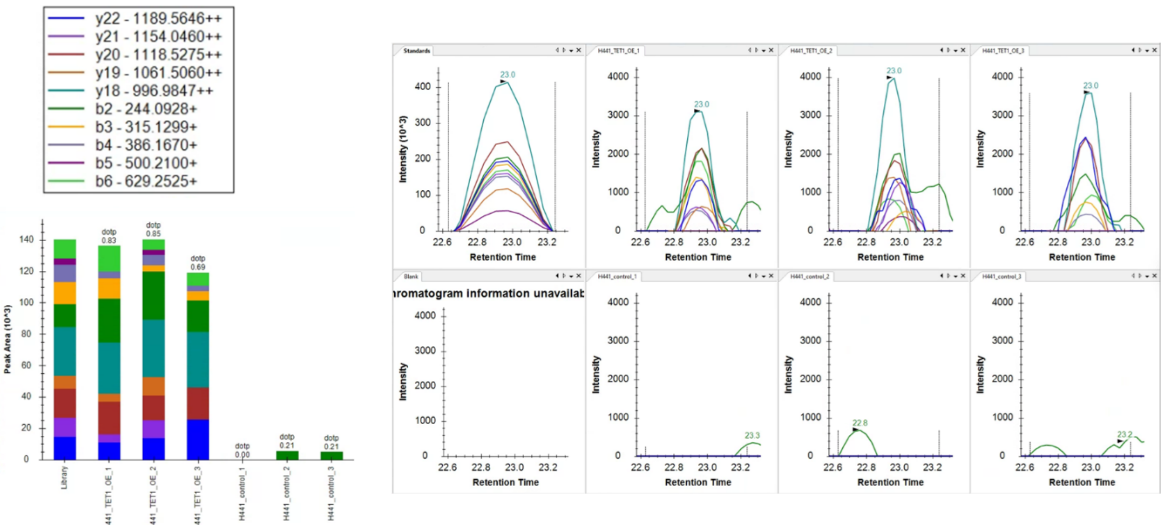  |
| b) |
| **TET1 Peptide: SLGVIPQDEQLHVLPLYK, 683.7175+++** |

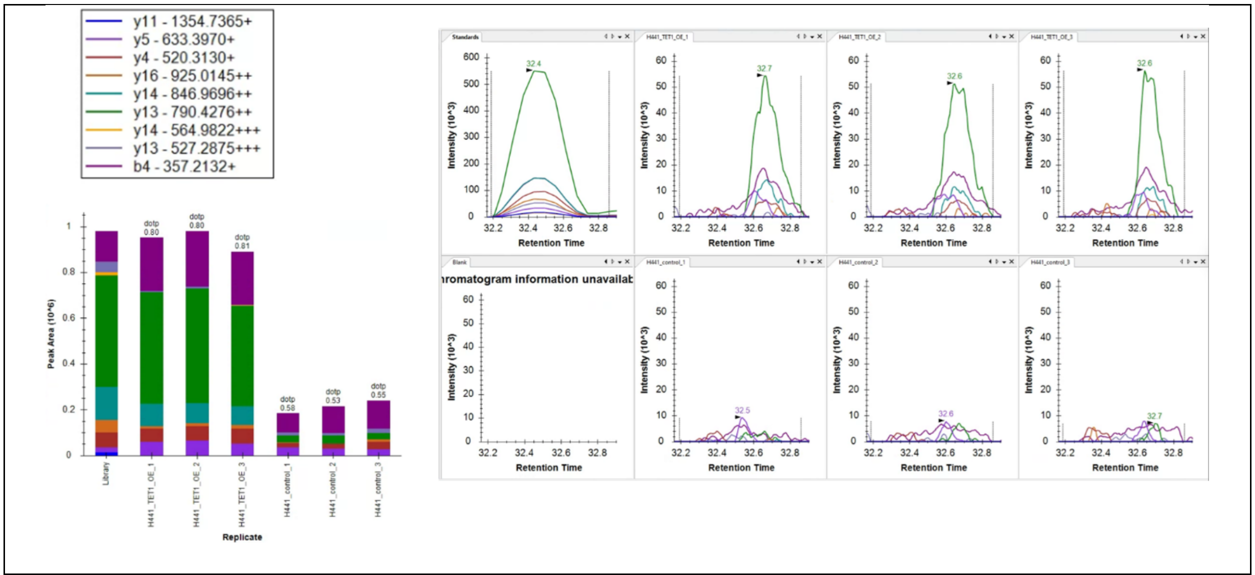


c)

**TET1 Peptide: NLEDNLQSLATR, 687.3546++**

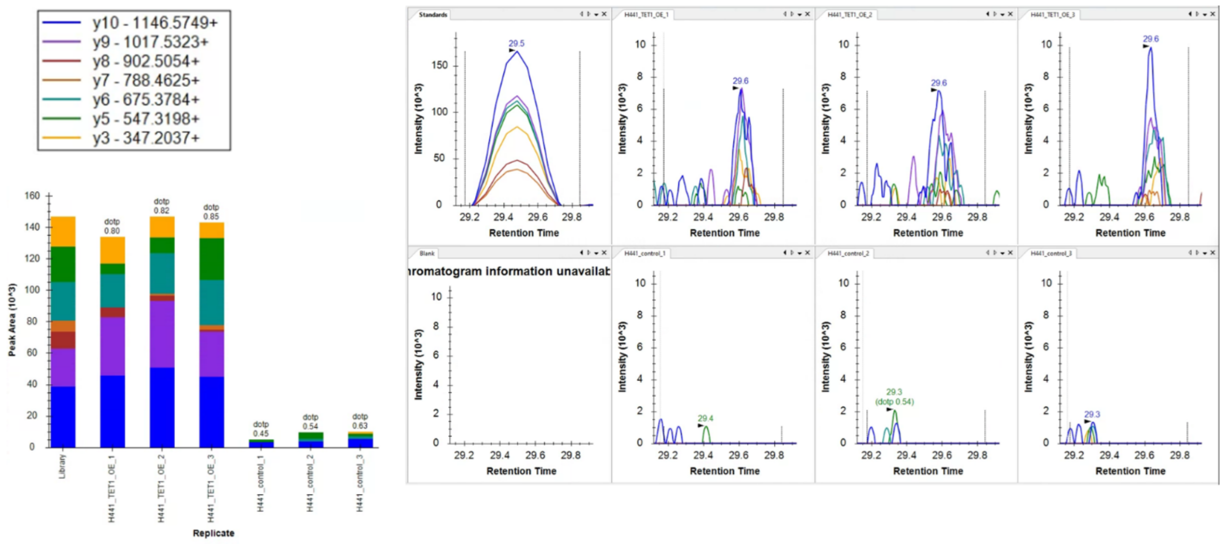


d)

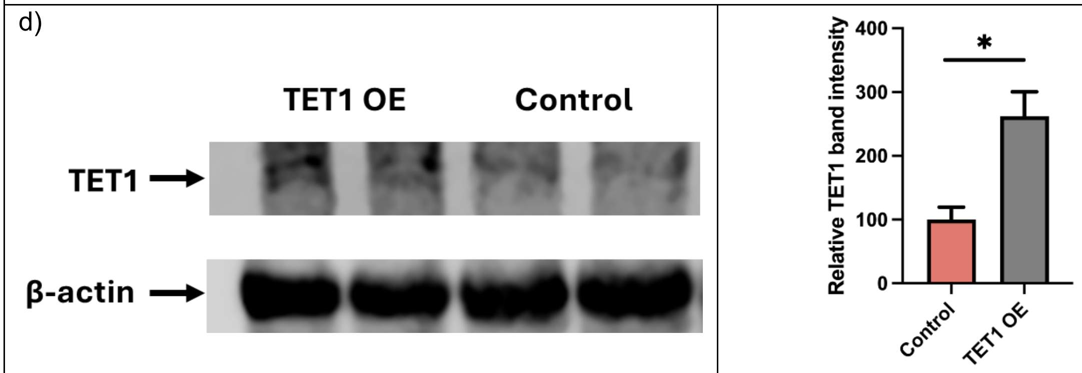

**Figure 2. LC-MS and western blotting characterization of TET1 protein in TET1 overexpressing H441 cells. a-c,** Three most intense TET1 peptide peaks showing various b and y ions. [Top extreme left panel is from TET1 standard peptide peak. Next top three panels are from TET1 overexpression sample. Bottom left panel is blank showing no peaks. And next bottom three panels are from control samples.] **d,** Left panel showing TET1 western blot in TET1 overexpression samples. The right panel is the corresponding bar diagram of TET1 band intensity from western blotting.

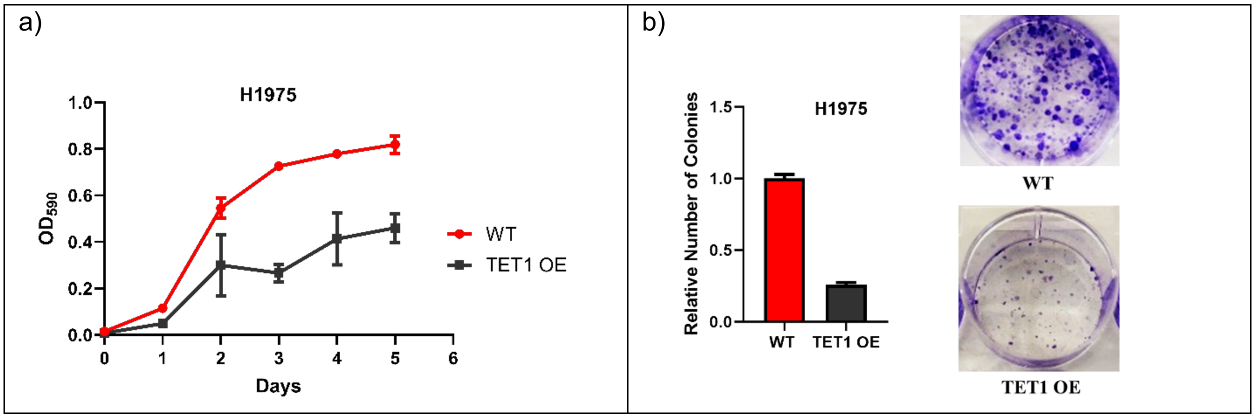


**Figure 3. Characterization of TET1 OE in H1975 NSCLC and downstream functional assays**. **a,** TET1 OE in H1975 cell lines lead to decrease in cell proliferation. **b,** Colony formation assay upon TET1 OE in H1975 cells. TET1 OE leads to a decrease in colony formation.

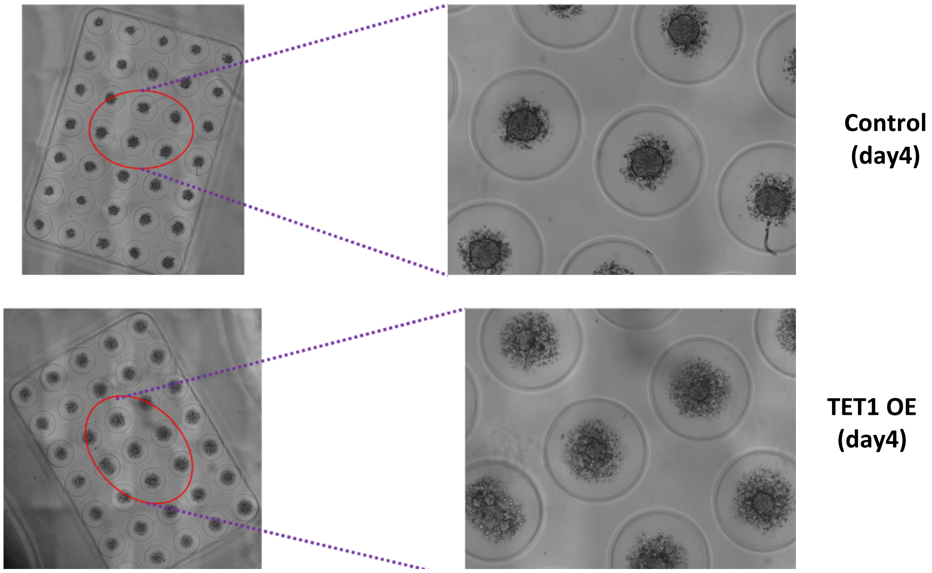


**Figure 4.** Pictorial representation of 3D spheroid image upon *TET1* overexpression in H441 cells.

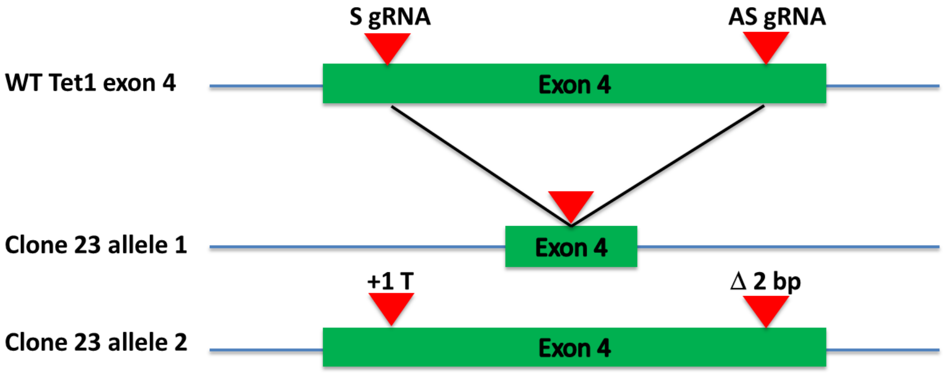


**Figure 5.** Illustration of the TET1 edited exon 4 for clone 23.

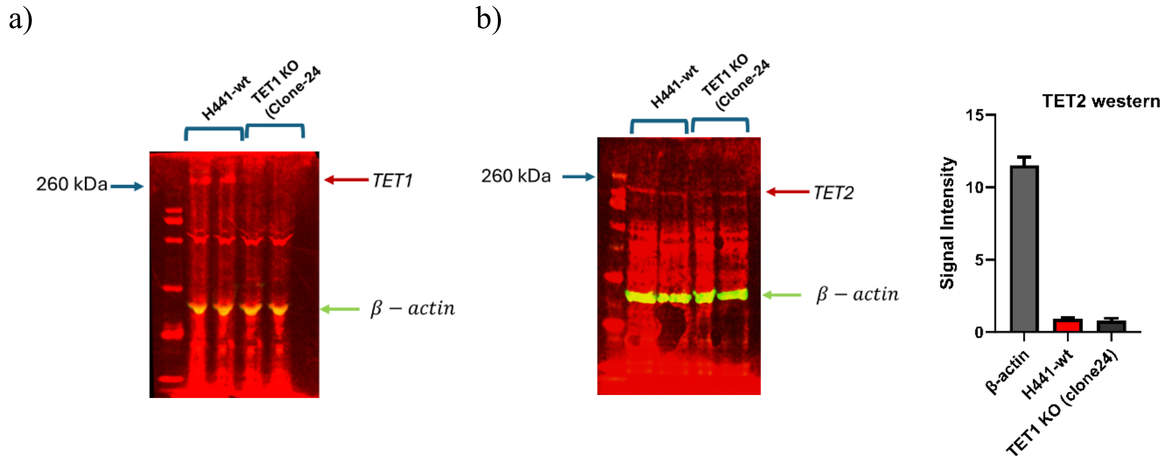


**Figure 6. Western blotting (WB) analysis of TET proteins in *TET1* KO clone. a,** WB analysis of TET1 in H441-WT and KO clone (Clone-24). **b,** WB analysis of TET2 in H441-WT and KO clone (clone-24).

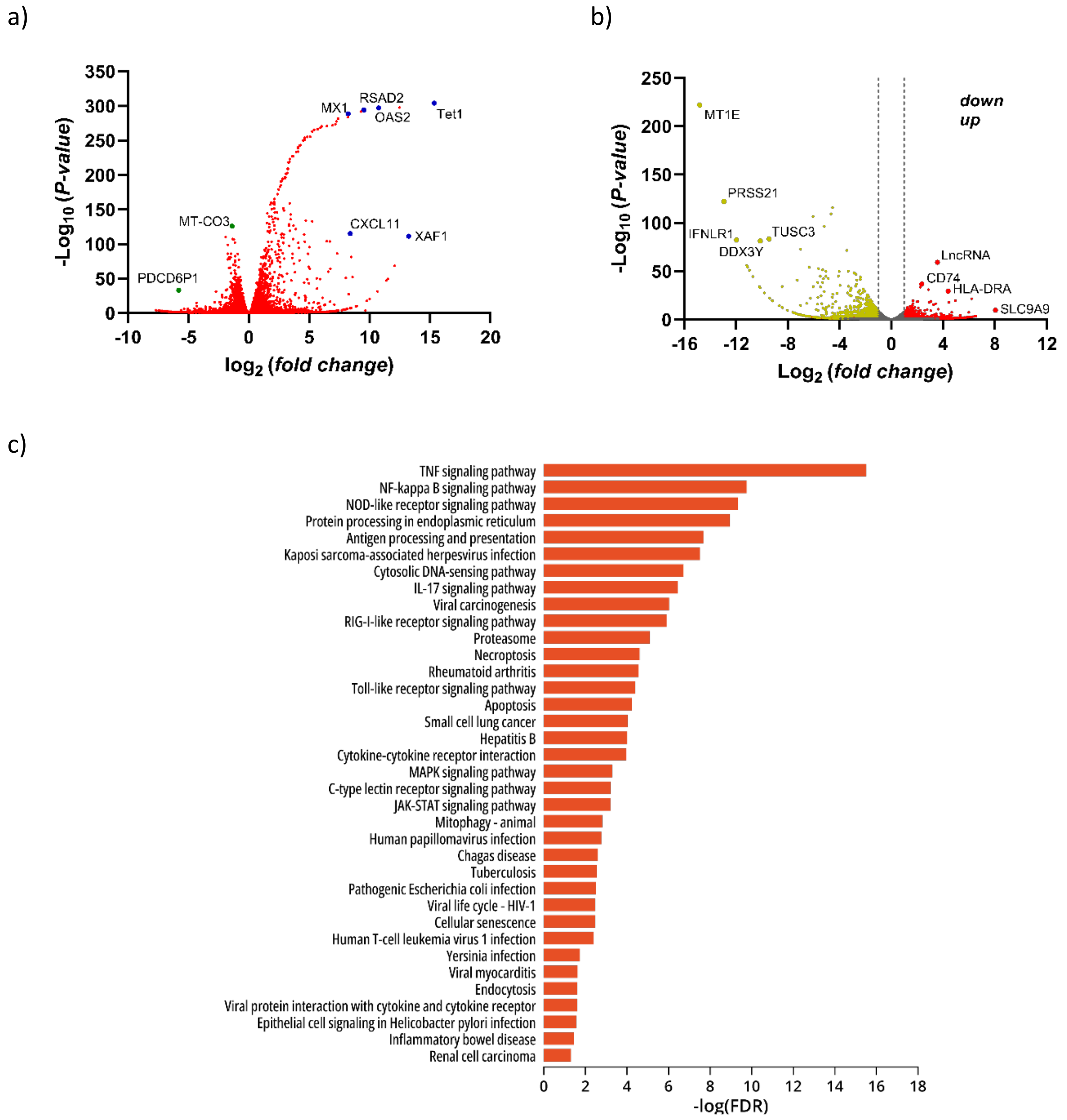

a)
350
300
250
200
150
100
50
0
-Log10 (P-value)
-10
-5
0
5
10
15
20
log2 (fold change)
MX1
RSAD2
OAS2
Tet1
MT-CO3
CXCL11
XAF1
PDCD6P1
b)
250
200
150
100
50
0
-Log10 (P-value)
-16
-12
-8
-4
0
4
8
12
Log2 (fold change)
down
up
MT1E
PRSS21
IFNLR1
TUSC3
DDX3Y
LncRNA
CD74
HLA-DRA
SLC9A9
c)
TNF signaling pathway
NF-kappa B signaling pathway
NOD-like receptor signaling pathway
Protein processing in endoplasmic reticulum
Antigen processing and presentation
Kaposi sarcoma-associated herpesvirus infection
Cytosolic DNA-sensing pathway
IL-17 signaling pathway
Viral carcinogenesis
RIG-I-like receptor signaling pathway
Proteasome
Necroptosis
Rheumatoid arthritis
Toll-like receptor signaling pathway
Apoptosis
Small cell lung cancer
Hepatitis B
Cytokine-cytokine receptor interaction
MAPK signaling pathway
C-type lectin receptor signaling pathway
JAK-STAT signaling pathway
Mitophagy - animal
Human papillomavirus infection
Chagas disease
Tuberculosis
Pathogenic Escherichia coli infection
Viral life cycle - HIV-1
Cellular senescence
Human T-cell leukemia virus 1 infection
Yersinia infection
Viral myocarditis
Endocytosis
Viral protein interaction with cytokine and cytokine receptor
Epithelial cell signaling in Helicobacter pylori infection
Inflammatory bowel disease
Renal cell carcinoma
0
2
4
6
8
10
12
14
16
18
-log(FDR)

d)

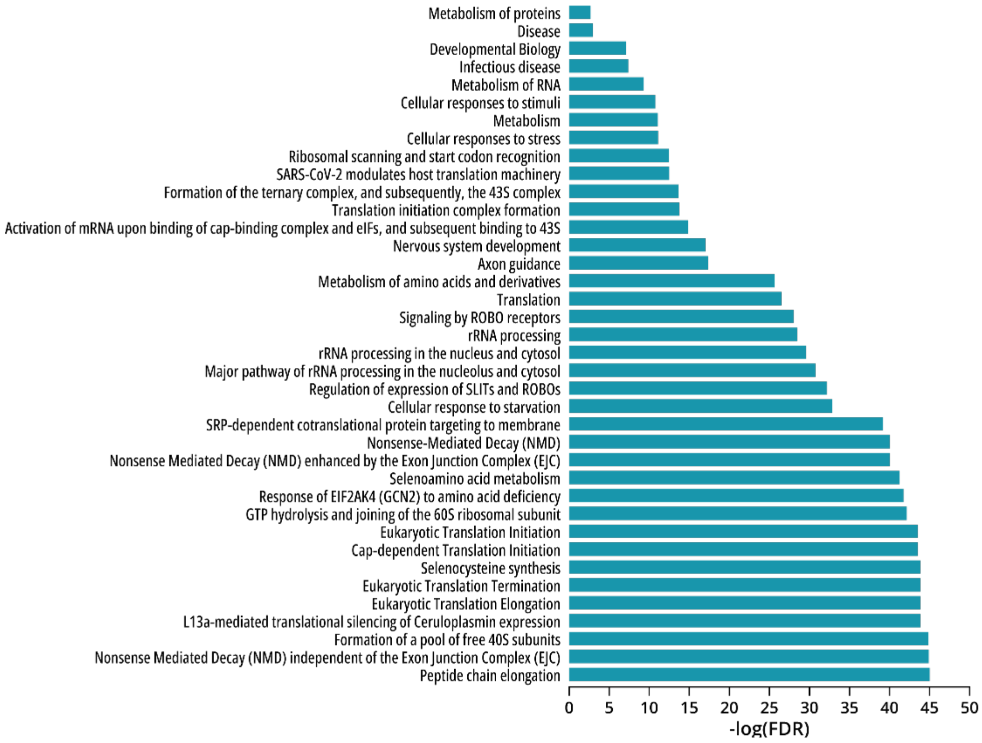


**Figure 7. Transcriptomics analysis upon *TET1* OE and KO in H441 cells. a,** Volcano plot showing differentially expressed genes upon *TET1* OE. **b,** Volcano plot showing differentially expressed genes upon *TET1* KO. **c,** KEEG biological pathways of significantly enriched upregulated genes upon *TET1* OE in H441 cells. **d,** KEEG biological pathways of significantly enriched upregulated genes upon *TET1* KO in H441 cells.

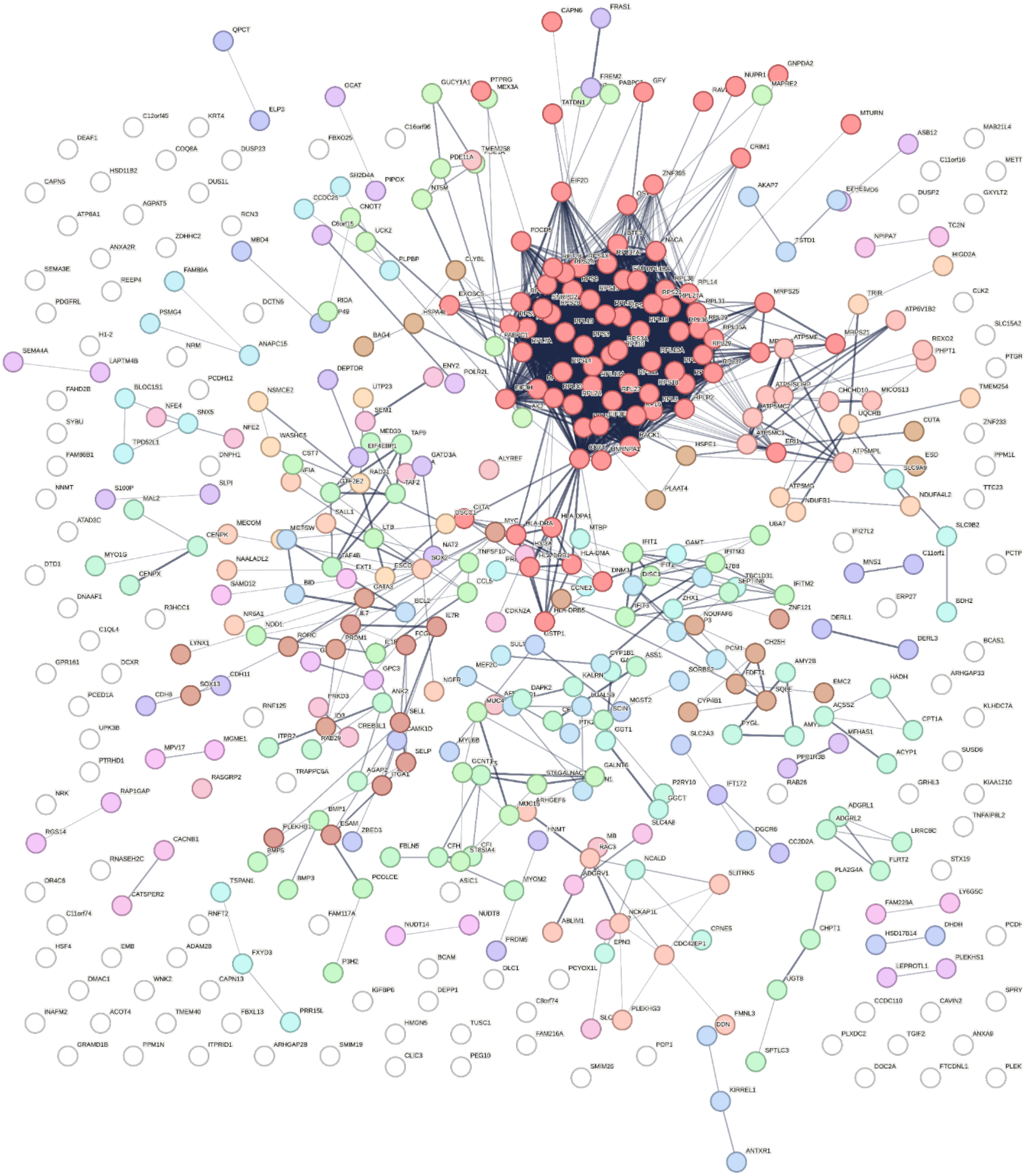

**Figure 8.** Upregulated gene interactome in *TET1* KO H441 cells (Clone 24).

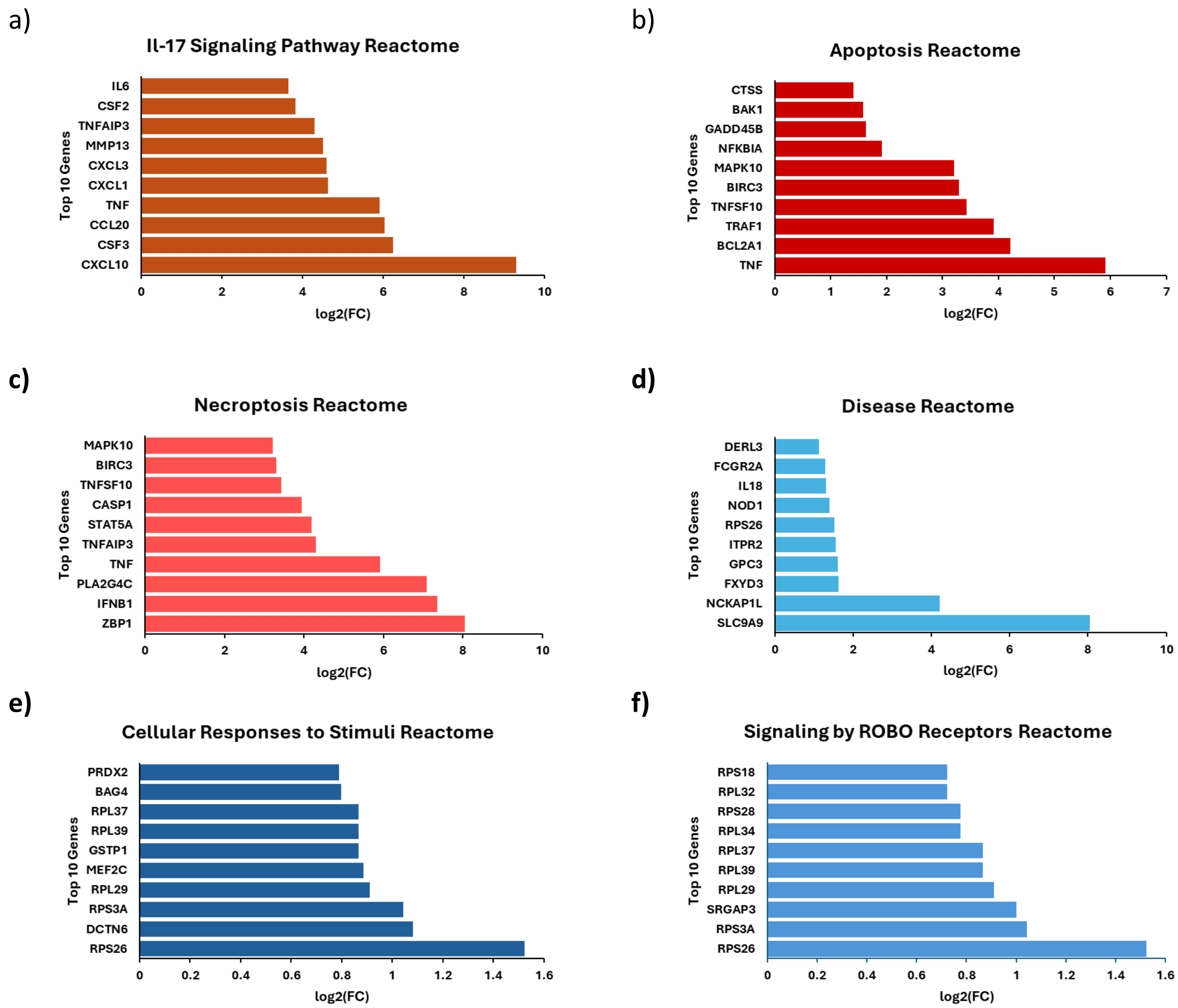


**Figure 9. Gene enrichment analysis on upregulated KEEG pathways in *TET1* OE and KO H441 cells. a-c,** Top 10 enriched genes in various upregulated signaling pathways upon *TET1* OE in H441 cells. **d-f,** Top 10 enriched genes in various upregulated signaling pathways upon *TET1* KO in H441 cells (clone 24).

a)

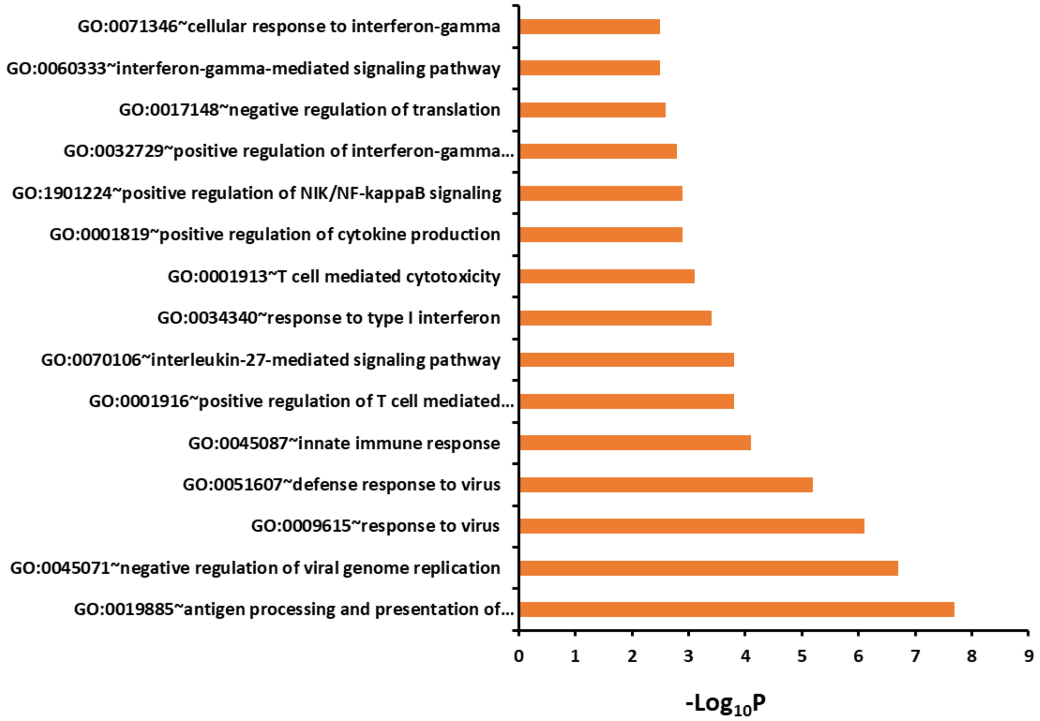


b)

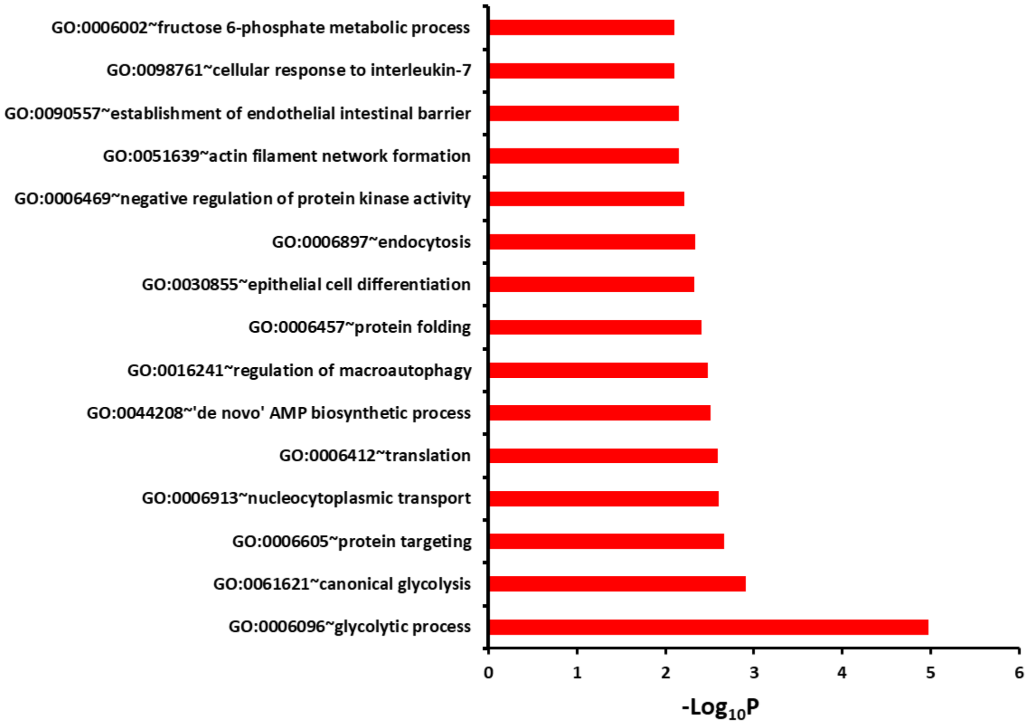


**Figure 10.** Top 15 GO terms for the biological processes in **a,** TET1 OE H441 cells and **b,** TET1 KO H441 cells (Clone 24).
